# Using singleton densities to detect recent selection in *Bos taurus*

**DOI:** 10.1101/2020.05.14.091009

**Authors:** Matthew Hartfield, Nina Aagaard Poulsen, Bernt Guldbrandtsen, Thomas Bataillon

**Affiliations:** Bioinformatics Research Centre, Aarhus University, DK–8000 Aarhus, Denmark; Institute of Evolutionary Biology, University of Edinburgh, Edinburgh EH9 3FL, UK; Department of Food Science, Aarhus University, Agro Food Park 48, 8200 Aarhus N, Denmark; Center for Quantitative Genetics and Genomics, Department of Molecular Biology and Genetics, Aarhus University, DK–8830 Tjele, Denmark; Rheinische Friedrich-Wilhelms-Universität Bonn, Institut für Tierwissenschaften, Tierzucht, Endenicher Allee 15, D-53115 Bonn, Germany; Department of Veterinary Sciences, Copenhagen University, DK–1870 Frederiksberg C, Denmark

**Keywords:** selection, genomics, *Bos taurus*, milk protein, milk fat, stature

## Abstract

Many quantitative traits are subject to polygenic selection, where several genomic regions undergo small, simultaneous changes in allele frequency that collectively alter a phenotype. The widespread availability of genome data, along with novel statistical techniques, has made it easier to detect these changes. We apply one such method, the ‘Singleton Density Score’, to the Holstein breed of *Bos taurus* to detect recent selection (arising up to around 740 years ago). We identify several genes as candidates for targets of recent selection, including some relating to cell regulation, catabolic processes, neural-cell adhesion and immunity. We do not find strong evidence that three traits that are important to humans – milk protein content, milk fat content, and stature – have been subject to directional selection. Simulations demonstrate that since *B. taurus* recently experienced a population bottleneck, singletons are depleted so the power of SDS methods are reduced. These results inform on which genes underlie recent genetic change in *B. taurus*, while providing information on how polygenic selection can be best investigated in future studies.

**Impact statement:** Many traits of ecological or economic importance (including height, disease propensity, climatic adaptation) are ‘polygenic’. That is, they are affected by a large number of genetic variants, with each one only making a small contribution to a trait, but collectively influence variation. As selection acts on all of these variants simultaneously, it only changes the frequency of each one by a small amount, making it hard to detect such selection from genome data. This situation has changed in recent years, with the proliferation of whole–genome data from many individuals, along with the development of methods to detect the subtle effects of polygenic selection. Here, we use data from 102 genomes from domesticated cattle (*Bos taurus*) that has experienced intense artificial selection since domestication, and test whether we can detect signatures of recent selection (arising up to 740 years ago). Domesticated species are appealing for this kind of study, as they are subject to extensive genome sequencing studies, and genetic variants can be related to traits under selection. We carried out our analysis in two parts. We first performed a genome–wide scan to find individual genetic regions that show signatures of recent selection. We identify some relating to cell regulation, catabolic processes, neural-cell adhesion and immunity. In the second part, we then analysed genetic regions associated with three key traits: milk protein content, milk fat content, and stature. We tested whether these regions collectively showed a signature of selection, but did not find a significant result in either case. Simulations suggest that the domestication history of cattle affected the power of these methods. We end with a discussion on how to best detect polygenic selection in future studies.

## Introduction

Determining which genomic regions have been subject to selection is a major research goal in evolutionary genetics. Traditional methods have focused on detecting strong selection affecting individual genes (Nielsen, 2005; Vitti *et al.*, 2013; Stephan, 2019). An alternative process is ‘polygenic selection’, where many loci contribute to genetic variation in a trait, so selection acting on it is expected to generate small and simultaneous allele frequency changes at multiple loci (Pritchard & Di Rienzo, 2010; Pritchard *et al.*, 2010). Many polygenic models have been formulated to account for both the response to phenotypic selection, and the maintenance of genetic variance in quantitative traits [reviewed by Sella & Barton (2019); Barghi *et al.* (2020)]. Among them is Fisher’s infinitesimal model, which is important for its historical role in uniting population and quantitative genetics, and its recent renaissance in the context of genome–wide association studies (Fisher, 1918; Barton & Keightley, 2002; Barton *et al.*, 2017; Charlesworth & Edwards, 2018; Visscher & Goddard, 2019). However, whereas it has been possible to identify which genetic regions contribute to trait variation, it has historically been hard to infer which alleles have been involved in the polygenic selection response. Extensive theoretical studies of how alleles at multiple loci act when a population adapts to a new optimum generally find that ‘large–effect’ alleles, which strongly affect a trait, are the first to spread and fix while ‘small–effect’ alleles take much longer to reach high frequencies (de Vladar & Barton, 2014; Wollstein & Stephan, 2014; Jain & Stephan, 2015, 2017a, 2017b; Stetter *et al.*, 2018; Thornton, 2019; Hayward & Sella, 2019). Furthermore, if epistasis exists between variants, many selected alleles do not reach fixation as they eventually become deleterious (de Vladar & Barton, 2014; Jain & Stephan, 2017b). The spread of large–effect alleles may also be impeded if a faster adaptive response can be otherwise realised through changes at many small–effect alleles (Lande, 1983; Chevin & Hospital, 2008; Pavlidis *et al.*, 2012; Chevin, 2019). Alternatively, if the optimum shift is sufficiently big, then large-effect mutations that first go to fixation can subsequently be replaced by small–effect variants over longer timescales (on the order of the population size; Hayward and Sella (2019)). Overall, only a small proportion of loci affected by polygenic selection are expected to fix sufficiently quickly to leave selection signatures in genomic data (Pavlidis *et al.*, 2012; Thornton, 2019).

Due to this difficulty, earlier methods for detecting polygenic selection focused on cases where selection favours distinct phenotypes in different populations, so trait differentiation amongst populations will be greater than expected under neutral drift. Tests for this form of selection relied on comparing *Q*_*st*_ and *F*_*st*_ statistics, which respectively measured mean genetic differentiation at the trait itself and a set of neutral loci (Whitlock, 2008; Le Corre & Kremer, 2012; Savolainen *et al.*, 2013). Yet these methods do not determine which genomic regions are subject to selection. This situation has now changed with the increased number of genome–wide association study (GWAS) data that link genotypes and phenotypes, as exemplified by the development of large cohort studies [e.g., the UK Biobank; Bycroft *et al.* (2018)]. The release of these data spurred a series of studies and new methods designed specifically to detect polygenic selection. These methods usually involve determining, which SNPs affecting a phenotype show correlated changes in frequency (Berg & Coop, 2014; Racimo *et al.*, 2018; Sanjak *et al.*, 2018; Josephs *et al.*, 2019; Berg *et al.*, 2019a, 2019b; Uricchio *et al.*, 2019; Edge & Coop, 2019; Kreiner *et al.*, 2020; Wieters *et al.*, 2021; Gramlich *et al.*, 2021); which sets of alleles are associated with certain environmental or climatic variations (Coop *et al.*, 2010; Turchin *et al.*, 2012; Robinson *et al.*, 2015; Yeaman *et al.*, 2016; Exposito-Alonso *et al.*, 2018; Zan & Carlborg, 2018; Exposito-Alonso *et al.*, 2019; MacLachlan *et al.*, 2021; Ehrlich *et al.*, 2021; Fuhrmann *et al.*, 2021; Rowan *et al.*, 2021); or determining which SNPs or genetic regions explain a large fraction of phenotypic variance and trait heritability (Zhou *et al.*, 2013; Yang *et al.*, 2015; Gazal *et al.*, 2017; Zeng *et al.*, 2018; Schoech *et al.*, 2019; Exposito-Alonso *et al.*, 2020; Duntsch *et al.*, 2020; Zeng *et al.*, 2021). Some of these approaches use overlapping methods.

Detecting recent polygenic selection is much harder, as long periods of time (number of generations on the order of the population size; Hayward and Sella, 2019; Thornton, 2019) may be needed to cause detectable frequency changes in alleles with small effect sizes. Over shorter timescales, these frequency changes are expected to be more modest and harder to detect (Stephan, 2016; Jain & Stephan, 2017a). A recent breakthrough in detecting these subtle changes was the development of the ‘Singleton Density Score’ (SDS), a statistic tailored to detect recent and coordinated allele frequency changes over many SNPs (Field *et al.*, 2016). Recent selection at a locus favouring one variant will lead to a reduction in the number of singletons (i.e., variants that are only observed once) around it. The SDS detects regions that exhibit a reduction in the density of singletons, to determine candidate regions that have been subject to recent selection. Using this approach, Field et al. (2016) found correlations between SDS scores at SNPs and their associated GWAS effect sizes for several polygenic traits in the modern UK human population, including increased height, infant head circumference and fasting insulin. Their findings suggested that these traits have been subject to recent selection during the most recent 75 or so generations (about 2,000 years). However, these (and other) results that detect selection for increased height may instead reflect previously unaccounted–for population structure (Novembre and Barton, 2018; Barton et al., 2019; Sohail et al., 2019; Berg et al., 2019; Uricchio et al., 2019; Edge and Coop, 2019).

The SDS method is ideally suited to organisms where large amount of whole-genome data are available, along with QTL or GWAS information that link genotypes to phenotypes, Domesticated species are attractive systems for studying recent selection, as selected phenotypes are often already known and these species are subject to large–scale sequencing studies. Investigating the genetic architecture underlying rapid selection in these species is also important to determine how they respond to agricultural practices, and uncover selection targets that can be used to improve breeding programs (Georges *et al.*, 2018). Domestic cattle *Bos taurus* has been subject to intensive genomics analyses to improve artificial selection for traits that are important for human use, including milk protein yield, milk fat content, and stature (Hayes *et al.*, 2009; Meuwissen *et al.*, 2013; Wray *et al.*, 2019). These traits are influenced in part by an individual’s genome, with significant heritability estimates being recorded, some as high as 80% (Soyeurt *et al.*, 2007; Haile-Mariam *et al.*, 2013; Buitenhuis *et al.*, 2016). Previous selection scans on *B. taurus* reported individual regions that were likely to be subject to recent selection, some of which were close to genetic regions for stature and milk protein content (Lemay *et al.*, 2009; MacEachern *et al.*, 2009; Qanbari *et al.*, 2010; Boitard & Rocha, 2013; Qanbari *et al.*, 2014; Zhao *et al.*, 2015; Boitard *et al.*, 2016a; Bouwman *et al.*, 2018). However, stature and milk protein content are polygenic traits, with several genetic regions and QTLs associated with each (Lemay *et al.*, 2009; Boitard *et al.*, 2016a; Bouwman *et al.*, 2018; van den Berg *et al.*, 2020). While recent methods have been developed to detect polygenic environmental adaptation (Rowan *et al.,* 2021), there has yet to be a formal test of whether these intrinsic traits show evidence of polygenic selection. Here, we applied the SDS method to whole–autosome sequencing data from 102 *B. taurus* Holstein individuals. We first determined genetic regions that have been subject to recent directional selection, and subsequently tested whether evidence exists for recent selection acting on a set of QTLs underlying either milk protein content, milk fat content, or stature in this breed.

## Results

### Methods outline

We filtered the data to retain only bi–allelic SNPs that had a sensible level of coverage and did not lie in putatively over–assembled regions (i.e., duplicated sections that caused many reads to assemble at a specific genetic location). Over– assembled regions appear as highly heterozygous with elevated coverage, and can exhibit false signatures of recent selection. We also obtained a set of singletons and filtered them to retain high-quality variants where both alleles were equally well covered to remove potentially erroneous calls. We polarised test SNPs using outgroup sequences and applied the SDS test of Field et al. (2016) to detect recent selection, with increased SDS values reflecting selection favouring derived SNPs over ancestral variants. We standardised SDS scores with those of a similar frequency, so they are normally distributed [similar normalisation was also carried out by Field et al. (2016)]. These values are denoted sSDS for ‘standardised SDS’. Further details are available in the *Methods* in the Supplementary Text.

### Estimating timescale of selection

We first determined the timescale over which we expect to detect selection in *B. taurus* using the SDS method. SDS measures the changes in singleton numbers around putatively selected SNPs, relative to background numbers in the absence of selection. As singletons arise on the tips of the underlying gene trees, the average tip length in the genealogy of sequenced samples determines the timescale over which the SDS detects a signal (Field et al., 2016). As more haploid genomes are included in the study, the time to first coalescence between two samples decreases, reducing the tip lengths and therefore shortening the timescale over which SDS detects selection (Field *et al.*, 2016). We hence simulate tip-ages over a range of sample sizes to investigate how this timescale changes accordingly.

To calculate the mean tip age, we simulated gene genealogies under two scenarios. We first simulated the Holstein population demography inferred by Boitard et al. (2016b), which suggested that this population experienced a sudden decline in effective population size (*N*_*e*_) since domestication, but with a present–day *N*_*e*_ (∼793) that is much larger than that inferred from pedigree data [∼49; Sørensen et al. (2005)] or from temporal variation in SNP frequencies (∼48; Jiménez–Mena et al. 2016). Hence, we also simulated genealogies under a second model that used the Boitard et al. (2016b) demographic model, but with the present–day *N*_*e*_ set to 49. These scenarios will be referred to as the ‘High *N*_*0*_’ and ‘Low *N*_*0*_’ models, respectively.

Figure 1 shows simulation results. Depending on the assumed present–day *N*_*e*_, the tip length in our sample of 204 alleles (i.e., assuming two per diploid individual) goes back either 65 or 148 generations. Assuming 5 years per generation (Boitard *et al.*, 2016b), this timescale corresponds to between 325 and 740 years ago. Since *B. taurus* domestication started around 10,000 years ago (Zeder, 2008) the sample size used in this study will only capture selection acting in the very recent past that is more relevant for breed formation, rather than selection during *B. taurus* domestication. Sample sizes and tip-ages are linearly related on a log-log scale, meaning that an increase in sample size will greatly decrease the timescale over which SDS detects selection. For example, with 500 haplotypes then SDS will detect selection acting no more than 50 generations ago, depending on the underlying demographic model.

**Figure 1:**
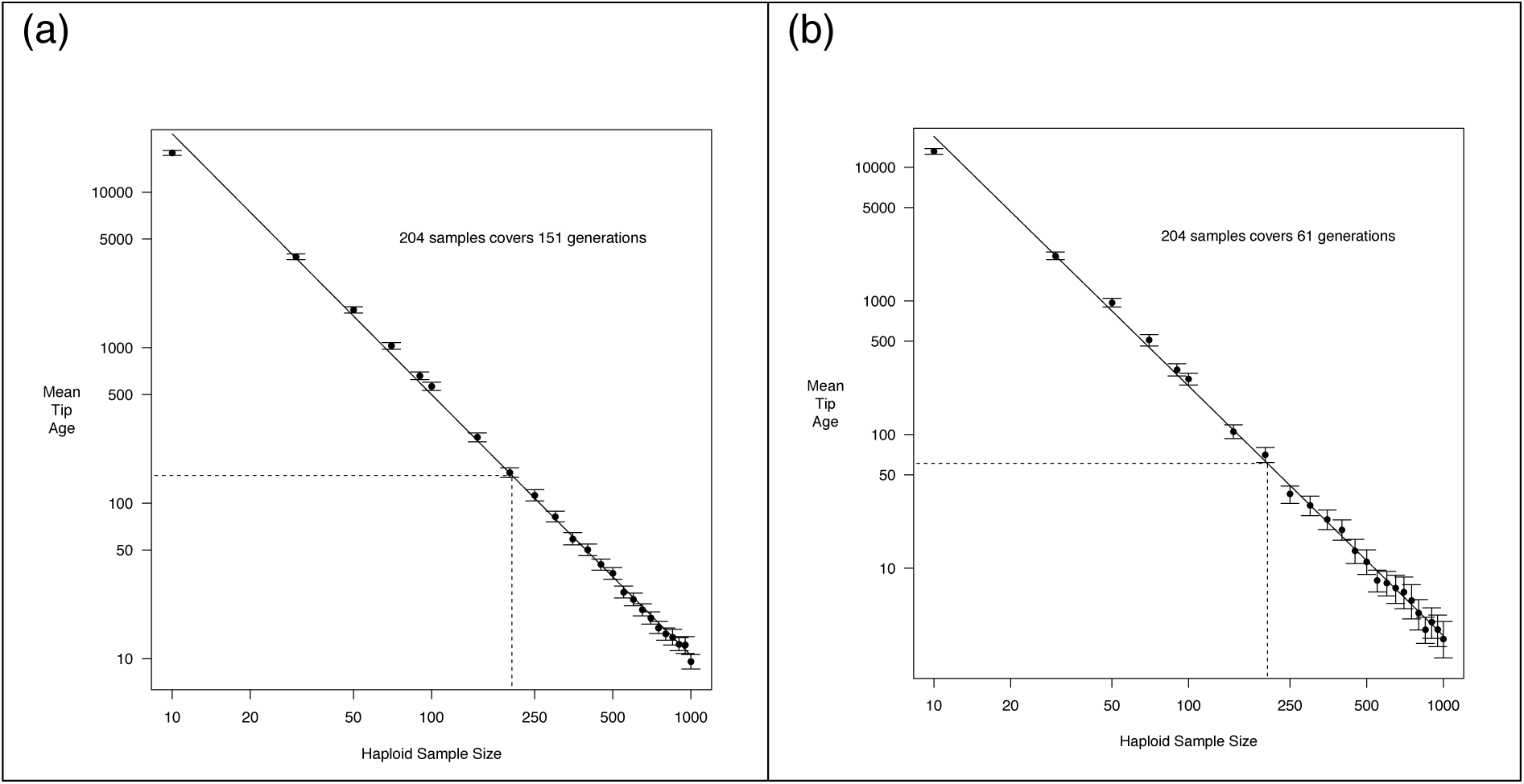
Simulated mean tip age for *B. taurus*, as a function of the number of haploid samples. Simulations assumed either (a) demography as inferred by Boitard et al. (2016b) (the ‘High *N*_0_’ model), or (b) the same but with a smaller present–day *N*_e_ of 49 (the ‘Low *N*_0_’ model). Points are the mean values; bars show 95% confidence intervals. The solid line is the best linear fit to the log of both values; dotted lines show the predicted tip age for 204 alleles.

We will focus on detecting selection signatures assuming the high *N*_*0*_ model. Results using the low *N*_*0*_ model to calibrate scores were broadly similar. They are outlined in the Supplementary Text; we will highlight when differences arise.

### Genome–wide sSDS

Figure 2 plots sSDS values (at SNPs with minor allele frequency greater than 5%) across all autosomes, excluding chromosome 25 (due to an insufficient number of singletons needed to obtain SDS scores after filtering). Many SNPs have elevated sSDS scores (158 SNPs at *FDR* < 0.05; 306 for the low *N*_*0*_ model). Several regions contain SNPs with significantly high sSDS values (Bonferroni–corrected nominal *P* < 0.05; actual *P* < ∼2.7 × 10^−8^). To further investigate potential selection targets, we looked for genes that either overlapped significant SNPs or lay 10kb up– or downstream of them. Linkage disequilibrium (LD), as measured by *r*^*2*^, decays to around 0.2 over 50kb in Danish Holstein breeds (Buitenhuis *et al.*, 2016), so genes within 10kb should be in LD with regions harbouring high sSDS scores. Table 1 lists these genes, with more targets present under the low *N*_*0*_ model. Most of these genes are of unknown function (as listed on UniProt); the list also includes an snRNA. *PPM1L* is involved with cellular regulation and the activation of stress–activated protein kinases. *TDO2* is involved in tryptophan–related catabolic processes, while *NTM* is implicated in neural cell adhesion. SNPs with significantly elevated scores are also found on chromosome 23 near the MHC region, which may reflect over– dominant selection. All Bonferroni–significant SNPs were removed from subsequent tests of recent polygenic selection to prevent directional selection from skewing the underlying sSDS distributions. Figure S1 shows results for the low *N*_*0*_ model.

**Figure 2:**
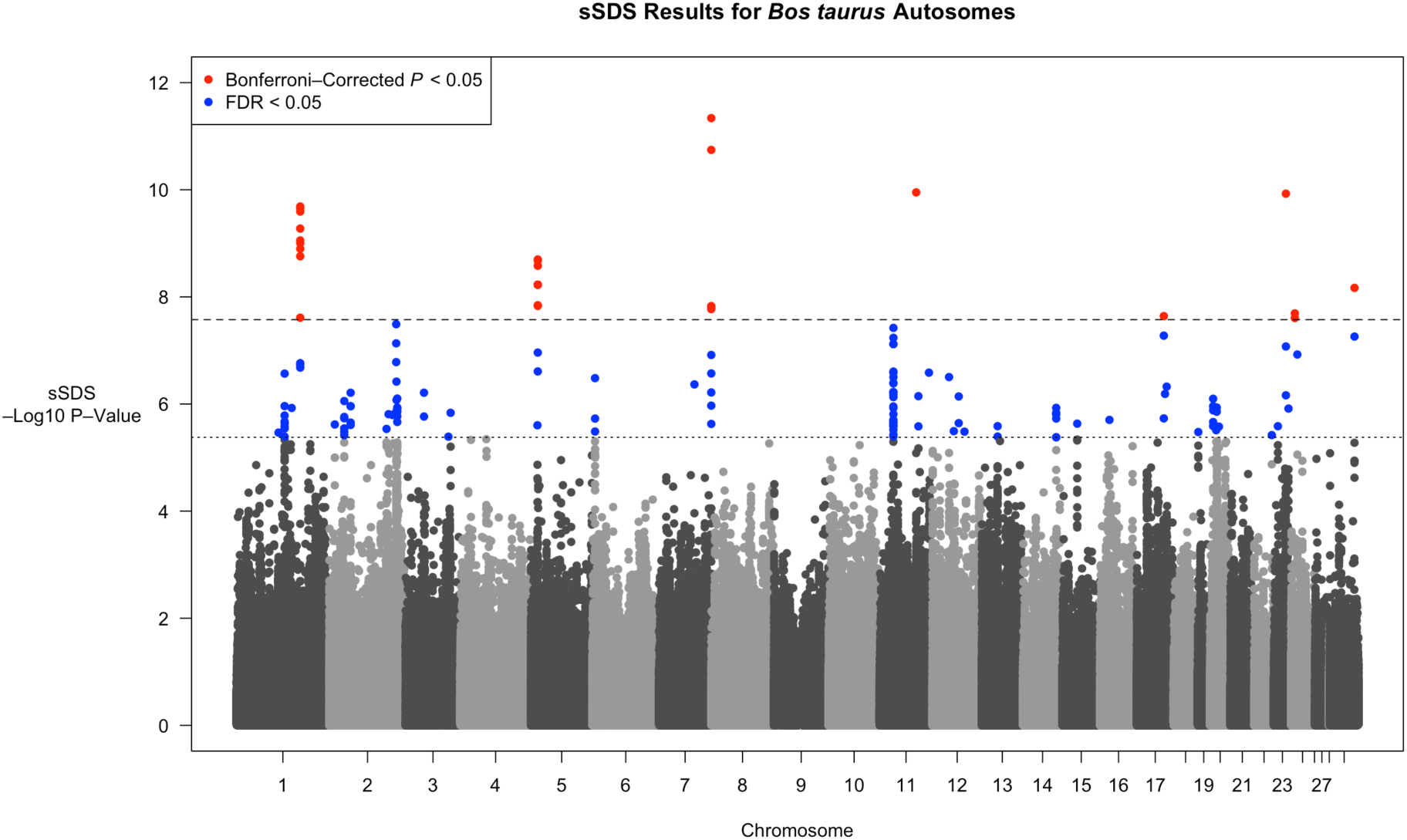
*P*–values of sSDS scores across *B. taurus* autosomes, as plotted on a negative Log_10_ scale, as a function of the chromosome. Alternating black and grey points show (non–significant) values from different chromosomes. Blue points are SNPs with *FDR* < 0.05, with the cut-off denoted by a horizontal dotted line. Red points are SNPs with Bonferroni–corrected *P*–value < 0.05 (actual *P*–value < ∼2.7 × 10^−8^), with the cut-off denoted by a horizontal dashed line. Figure S1 shows results for the Low *N*_0_ model.

**Table 1:** Genes that overlap or lie close to Bonferroni–significant sSDS regions. The ‘High, Low *N*_0_’ column specifies which genes are close to significant SNPs for each *N*_0_ model.

| Chromosome | Gene Name | Start Position | End Position | Gene Biotype | High, Low $N_0$ |
| --- | --- | --- | --- | --- | --- |
| 1 | PPM1L | 106405113 | 106727070 | Protein Coding | High, Low |
| 5 | TMCC3 | 24306913 | 24595494 | Protein Coding | High, Low |
| 5 | CEP83 | 24070404 | 24345243 | Protein Coding | High, Low |
| 17 | U6 | 43381106 | 43381209 | snRNA | Low |
| 17 | CTSO | 43364999 | 43381605 | Protein Coding | Low |
| 17 | TDO2 | 43386894 | 43403747 | Protein Coding | High, Low |
| 23 | OR12D2H | 29291787 | 29292713 | Protein Coding | High, Low |
| 23 | OR12D2E | 29305933 | 29309785 | Protein Coding | High, Low |
| 24 | GAREM1 | 24694637 | 24927333 | Protein Coding | High, Low |
| 29 | NTM | 34576918 | 34994005 | Protein Coding | High, Low |

#### Testing for polygenic selection acting on milk protein and stature

If polygenic selection were acting on specific traits, we expect a positive correlation between the effect size of variant underpinning it, and selection acting on it as measured by sSDS. We collated sSDS scores of SNPs that lie close to QTLs reported for either milk fat percentage, milk protein percentage (van den Berg *et al.*, 2020), or those that lie close to stature QTLs (Bouwman *et al.*, 2018). The latter were inferred from a meta–analysis of GWAS studies conducted in seven Holstein populations, but not every QTL had an effect size reported in each population. We hence investigated two overlapping consensus QTL sets, where an effect size was either reported in at least 6 of 7 populations (yielding 42 QTLs with sSDS scores associated with them), or where effect sizes were reported in at least 5 of 7 populations (58 QTLs had sSDS scores). We re-polarised SDS scores so that a positive score reflected a trait-increasing effect; we denote these values ‘tSDS’ following Field *et al.* (2016). We then determine if there was a positive correlation between the absolute log_10_-value of the QTL *P*-value (a proxy for the effect size) and tSDS.

Figure 3 shows the relationship between QTL *P*-values and tSDS for SNPs that lie close to QTLs. Although positive trends are observed, they all exhibit non-significant correlations (milk fat percentage Spearman ρ= 0.0990, *P* = 0.603; milk protein percentage Spearman ρ= 0.0354, *P* = 0.758; stature from 6 breeds Spearman ρ= −0.0739, *P* = 0.642; stature from 5 breeds Spearman ρ= −0.00966, *P* = 0.943). Relationships remain non-significant after removing an outlier point for the milk traits whose QTL has an extremely low *P*-value (Figure S2), and also under the low *N*_*0*_ model (Figure S3; see figure legends for regression *P*-values).

**Figure 3:**
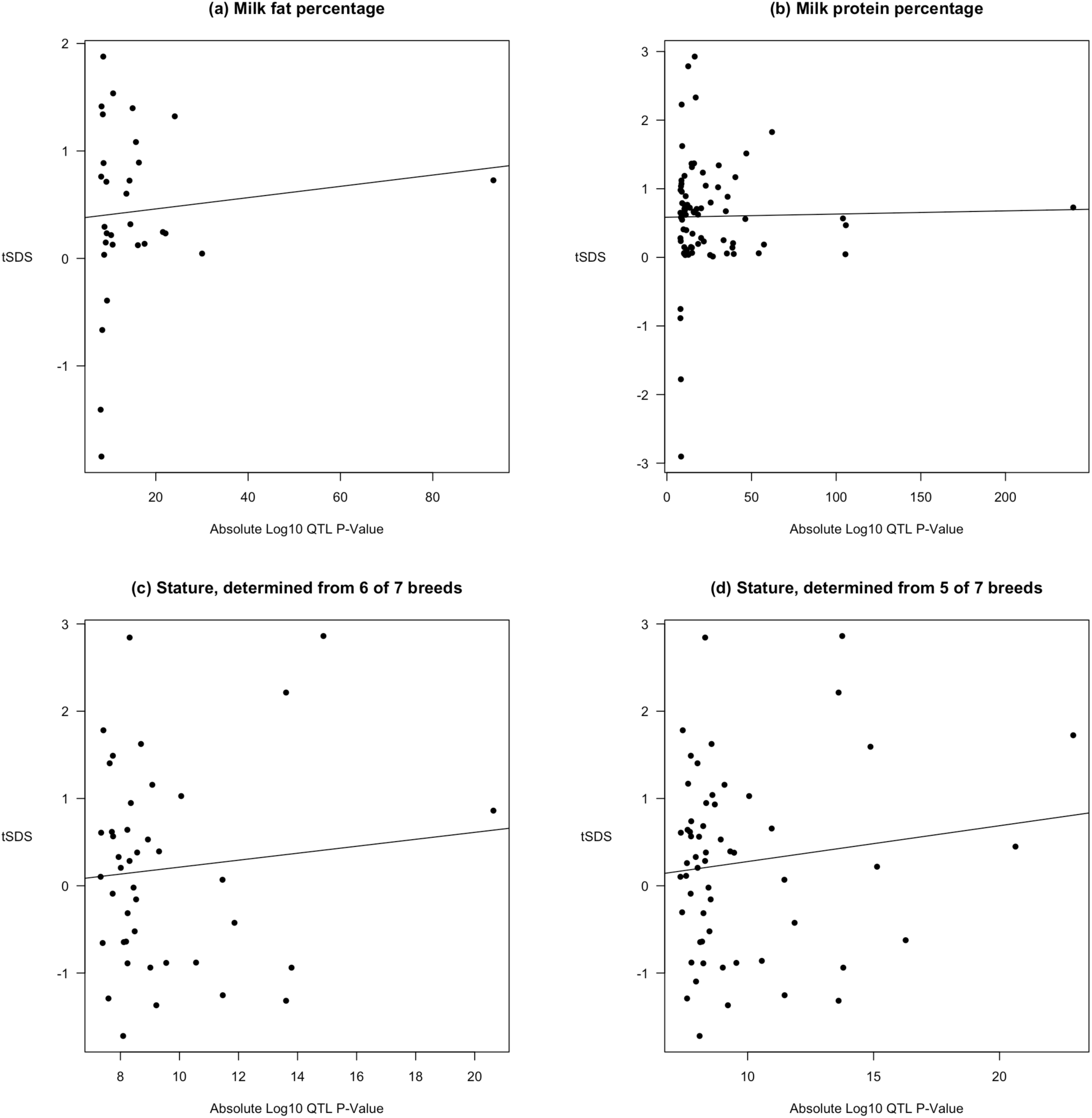
Relationship between tSDS scores near milk or stature QTLs, as noted in the subheadings, and the absolute log *P*-value of QTLs. Lines show a linear model regression fit. Figure S3 shows results assuming a low *N*_*0*_ model.

sSDS (and tSDS) can become correlated along the genome if focal SNPs are in LD with one another, which was not accounted for in the preceding analyses. To determine whether LD could have affected these correlations, we randomly subsampled sSDS scores from SNPs that shared the same chromosome and bin of derived-allele frequency as the SNPs used in the above analyses, and re-polarised them to transform them into tSDS values. We then determined the Spearman’s ρ associated with these permuted values to determine whether that for the true data was significantly elevated (see Methods for details). In all cases, the observed value was not significantly higher than for permuted values (see Figure S4 for histograms and exact *P*–values, which all exceed 0.05). We therefore conclude that these QTL datasets do not harbour SNPs with significantly different tSDS scores compared to the rest of the genome.

## Discussion

### Summary of results

We analysed an extensive *B. taurus* genomic dataset to identify signatures of recent selection in the Holstein breed, and to determine whether the data contained a signal of polygenic selection acting on milk proteins and QTLs underlying phenotypic variation in stature. Given the sample size and the demographic history of Holsteins, the SDS method can detect very recent selection events arising no more than approximately 740 years ago (Figure 1). A whole–genome scan for sSDS scores identified several targets of recent directional selection that overlap or lie close to protein–coding genes (Figure 2; Table 1). The genes whose functions are known are involved in protein regulation, catabolic processes, and neural-cell adhesion. Significant values were also observed near the MHC region. We subsequently investigated whether either milk protein genes or SNPs near stature QTLs collectively showed evidence of polygenic selection. We did so by testing whether there is a relationship between the QTL effect size, as measured by its *P*-value, and tSDS values to SNPs near them. However, no relationship was observed, even after performing a permutation test (Figures 3, S2-S4). Hence, while sSDS could reveal specific instances of recent selection, tests based on collective scores of variants associated with known selected traits yielded no signal of polygenic selection.

### Potential reasons for a lack of polygenic selection signal

#### Impact of Holstein demographic history

While the SDS method detected individual candidate genes for very recent selection, we were unable to find strong evidence for polygenic selection acting on three traits that were subject to artificial selection since domestication. This result is *a priori* surprising, given that these traits have been subject to recent intense artificial selection. Recent studies generally find non-zero heritability estimates for them, indicating that there should be the potential for genetic variants underpinning them to change in response to artificial selection (Soyeurt *et al.*, 2007; Haile-Mariam *et al.*, 2013; Buitenhuis *et al.*, 2016). In addition, the ratio of the mutation and recombination rates in cattle is just over three (Boitard *et al.*, 2016b; Harland *et al.*, 2018), indicating that several informative SNPs exist per haplotypes that should improve the power of the SDS method [in contrast, this ratio is approximately equal to one in humans (Field *et al.*, 2016)].

One potential reason for this lack of signal is due to the population history of *Bos taurus*. The effective population size of many *B. taurus* breeds appears to have undergone a decline since domestication (Sørensen et al., 2005; Boitard et al., 2016b), which likely reflects successive bottlenecks due to domestication, breed formation and intense recent selection. Population size reductions are known to reduce the number of low-frequency variants and increases the prevalence of intermediate-frequency variants (Harpending *et al.*, 1998), which can affect the power of the SDS method. To understand if the history of *B. taurus* affects the detection of recent selection in Holstein cattle using SDS, we ran coalescent simulations to determine its ability to detect ongoing selection, given realistic Holstein population history and genetic parameters (see Methods for details). We simulated a partial sweep occurring in the middle of a 10Mb region, either assuming a mutation rate in line with what has been inferred for Holstein, or one 10-fold higher to replicate diversity expected in a genetic region with an elevated mutation rate.

For the standard mutation rate, no SDS scores were produced for any simulations. After inspecting the simulation results, we see that there is a large skew in the distribution of singleton numbers per individual with a large number of individuals (over 20 on average) that do not carry singletons at the end of simulations, preventing the calculation of a local SDS score (Figure 4). This fraction remained the same irrespective of whether the simulated region was neutral or subject to selection; the main effect of a sweep was to reduce the mean number of singletons per individual, which is the signal measured by SDS (Field *et al.*, 2016). This reduction in overall singleton numbers is consistent with the known effects of population size contraction on reducing tip lengths (Harpending *et al.*, 1998).

**Figure 4:**
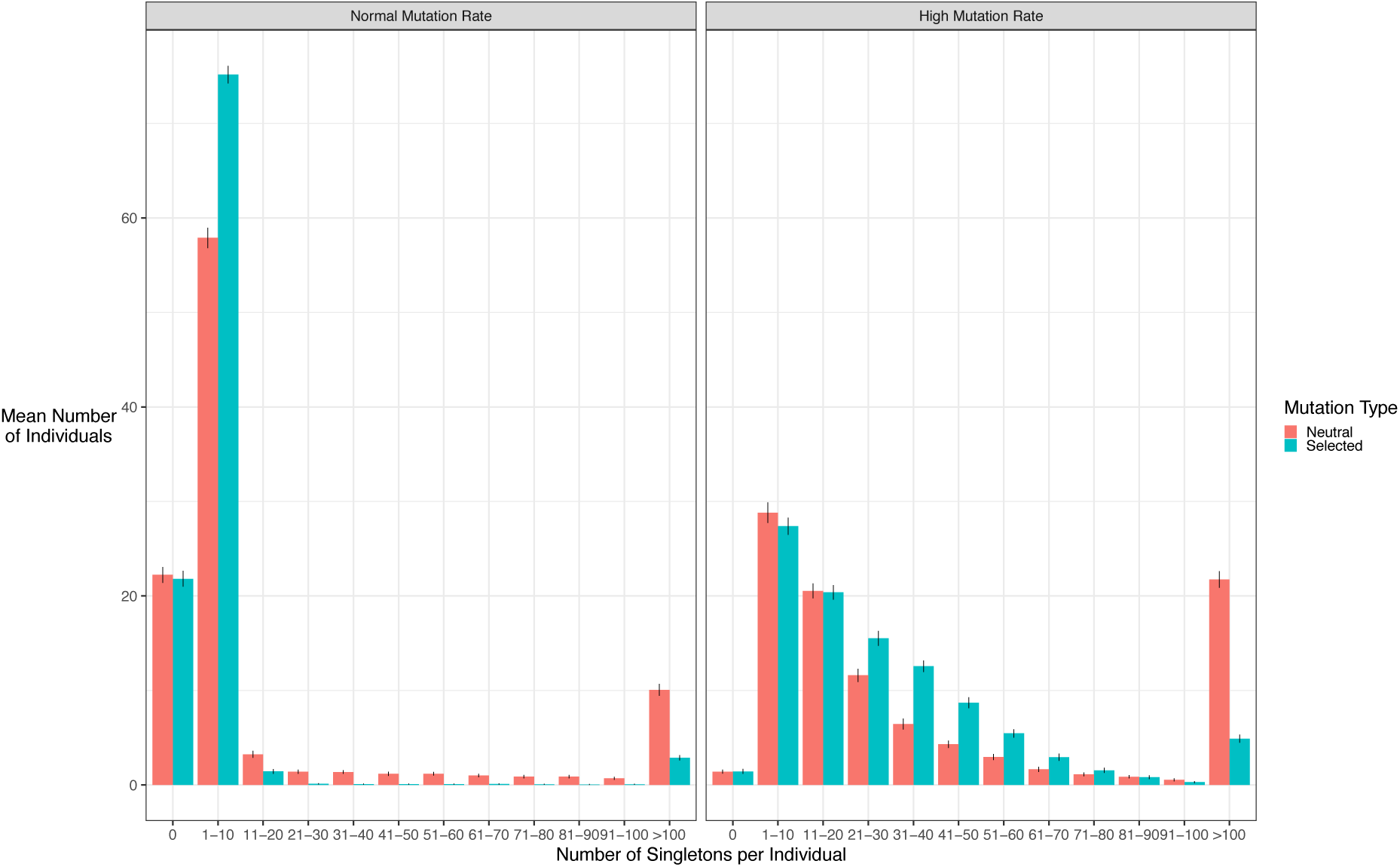
Mean distribution of singleton numbers per individual for each simulation, wither assuming a standard mutation rate (left) or a 10-fold higher mutation rate (right). Bars represent 95% confidence intervals.

With a 10-fold higher mutation rate, there were fewer cases where no individual harboured singletons (Figure 4). Accordingly, SDS scores could be calculated for 65 and 66 out of 100 simulations for the neutral and selective cases respectively. In these cases, sSDS values were significantly higher in the selected case than for the neutral case (Figure 5; two-sided Wilcox Test *P* = 1.1×10^−5^). However, note that sSDS values is less than one for the selected case, which does not exceed the FDR threshold in our study (for the high *N*_0_ case, the smallest sSDS value with FDR < 0.05 is 4.46).

**Figure 5:**
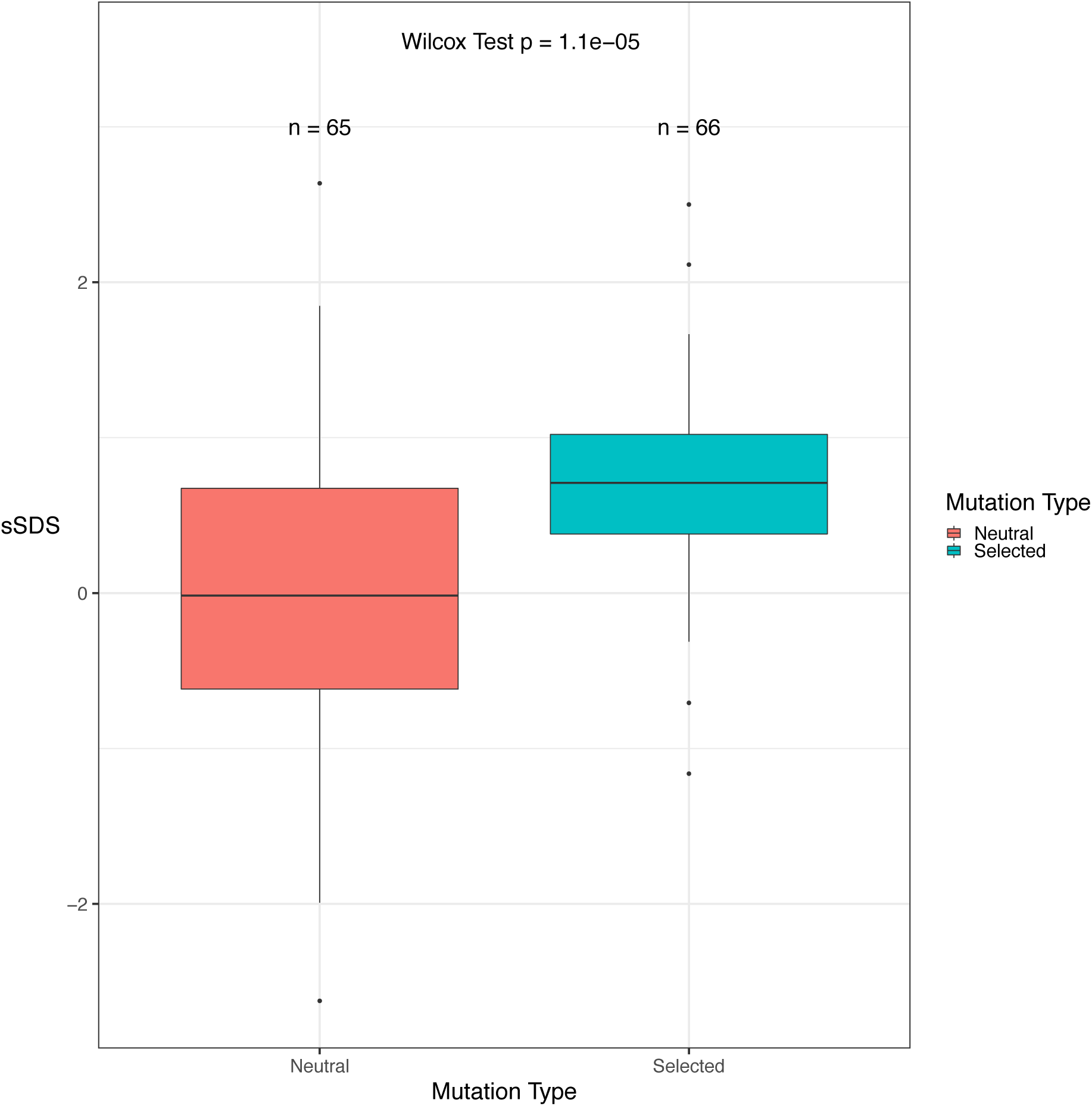
Distribution of simulated sSDS scores assuming a high mutation rate, for the neutral and selected cases. Numbers above each box plot denotes how many simulations produced SDS scores and were included in the plot.

Although singleton numbers differ between the two cases, a reduction in power could also be caused by a more general reduction in diversity due to the small recent effective population sizes of cattle. To investigate this effect, we estimated the fixed *Ne* that would yield the same number of segregating sites in simulations using the standard mutation rate, based on Watterson’s estimator (Watterson, 1975; Hudson, 1990; see Methods for details). In both cases where selection is present or absent, *Ne* estimates lie at around 25,000, which is that inferred at approximately halfway between the onset of domestication and the present day. (Boitard *et al.*, 2016b; Figure S5). Given that estimates are similar irrespective of whether a sweep was present or not, the reduced population size caused by domestication could have also affected power due to limiting genetic variation and thus the potential to detect subtle sweep signatures associated with polygenic selection.

Overall, these simulations are consistent with population size reductions in *B. taurus* both reducing the overall genetic diversity and the number of singletons, which limits its ability to detect partial sweeps. SDS is more likely to detect signals in regions of elevated mutation rate, suggesting there will likely be an ascertainment bias in where signals are detected in the genome. The reduction in singletons also reduces the power to investigate SDS values in telomeric regions. SDS values are calculated using the distance up- and downstream from a SNP to the nearest singleton, and are undefined if a certain number of samples do not harbour singletons in either direction (Field *et al.*, 2016). SDS values are hence less likely to be defined in telomeric regions, as it is generally less feasible to observe singletons up until the end of the chromosome. This problem is exacerbated if there are few singletons overall.

#### Other potential reasons for a lack of signal

Another potential reason for a lack of signal is that the selection response on these traits may have been driven by large-effect variants that have already fixed in the population, with a smaller contribution from small-effect mutations. Theoretical models have shown that more major–effect QTLs are likely to fix if the population lies further from a fitness optimum (Lande, 1983; Jain & Stephan, 2017b; Thornton, 2019). Domesticated species, which experience strong and sustained directional artificial selection, especially in recent generations, could thereby fix more adaptive mutation via sweep–like processes compared to populations evolving in more stable environments (Lande, 1983; Jain & Stephan, 2017a). Furthermore, once a population has adapted to a new environment (the domestication phenotype in this case), then any remaining major–effect mutations are likely to be superseded by variants with weaker effects, which are harder to detect (Hayward & Sella, 2019). The response to polygenic selection will be further weakened in smaller populations (John & Stephan, 2020), which could be a factor given the reduced effective population sizes of *B. taurus* (Sørensen *et al.*, 2005; Boitard *et al.*, 2016b). There is some evidence of this explanation; selective sweeps signatures are associated with stature QTLs (Bouwman *et al.*, 2018), and the study of van den Berg et al. (2020) was more likely to identify milk QTLs that had a moderate to high minor allele frequency, suggesting reduced power to detect low-frequency variants that are potential contributors to polygenic selection. Conversely, the stature meta-analysis by Bouwman et al. (2018) found significant SNPs that explained up to 13.8% of the variance in stature, which is similar to that explained by significant SNPs for human height (16%), which is a classic trait for polygenic selection studies. Hence, there may be sufficient polygenic SNPs present to test for polygenic selection, but the power will still be reduced due to the demographic history of Holstein cattle.

Potential solutions to increase power include increasing sample sizes; using alternative methods; or analysing different kinds of genome data to detect polygenic selection. Applying SDS to a larger sample size would increase the power to detect selection acting in the recent past [Figure 1; see also Field et al. (2016)], but overall power will still be limited by the tip-length of neutral genealogies. Recent developments in methodology involve directly inferring trees from genome data, and using these to identify subtle sweep signatures associated with trait variants (Edge & Coop, 2019; Speidel *et al.*, 2019; Stern *et al.*, 2021). These methods have greater power to detect weakly-selected mutations that may be segregating for longer than the tip-length of the population.

Another approach would be to look beyond sequence data and focus on gene networks [reviewed by Fagny & Austerlitz (2021)]. The recently–proposed ‘omnigenic’ model (Boyle *et al.*, 2017; Liu *et al.*, 2019) posits that variation in quantitative traits is principally affected by a plethora of ‘peripheral’ genes that indirectly affect them, rather than a limited set of ‘core’ genes that directly modify a trait. These numerous peripheral genes may exert their influence via regulatory effects (e.g., gene expression changes), but are also expected to be highly pleiotropic. Fully testing the omnigenic model will require larger datasets and novel experimental designs (Wray *et al.*, 2018). A recent example is from an experiment with *Drosophila melanogaster*, where gene knockouts that do not pass a GWAS significance threshold for pupal length still significantly affect it (Zhang *et al.*, 2021). There is also nascent evidence that gene regulation may underlie directional polygenic selection. Boitard et al. (2016a) found that some adaptive signatures of *B. taurus* are located in intergenic regions; regulatory changes were also proposed to guide polygenic selection in *Arabidopsis* (He *et al.*, 2016). Analyses of gene–sets associated with infection responses or immunity also found evidence for polygenic selection in humans and primates (Daub *et al.*, 2013, 2017; Svardal *et al.*, 2017). Immunity gene–sets might be exceptional cases, as they are more likely to contain genes subject to very strong selection (Castellano *et al.*, 2019). Further investigations using regulatory information and a broader range of gene–sets could be a promising approach to determine the impact of polygenic selection.

## Materials and Methods

Full methods are available in the Supplementary Text.

## Acknowledgements

We would like to thank Simon Boitard for sharing his results on *B. taurus* demographic inference; Simon Boitard, Jon Slate and two anonymous reviewers for providing feedback on the manuscript. MH is supported by a NERC Independent Research Fellowship (NE/R015686/1). NAP, BG and TB are funded by a synergistic research grant from the Faculty of Science and Technology, Aarhus University, Denmark. MH and TB also acknowledge financial support from the European Research Council under the European Union’s Seventh Framework Program (FP7/20072013, ERC Grant 311341). The authors declare no conflicts of interest.

## Author contribution

All authors contributed to the study design. NAP and BG provided data. MH performed the analyses and wrote the manuscript, with feedback from NAP, BG and TB.

## Data archiving

Raw SDS scores and polarisation information has been deposited on Dryad (https://doi.org/10.5061/dryad.547d7wm8q). Data analysis and simulation scripts are available on GitHub (https://github.com/MattHartfield/CattleSDS).

