## Supplementary Text for "Using singleton densities to detect recent selection in *Bos taurus*"

***Matthew Hartfield, Nina Aagaard Poulsen, Bernt Guldbrandtsen, Thomas Bataillon***

#### *Materials and Methods*

#### *Supplementary Figures*

**Figure S1:** Genome wide sSDS results for the low $N_0$ model.

**Figure S2:** Polygenic selection test results for the high $N_0$ model (milk fat percentage, milk protein percentage) after removing outlier point.

**Figure S3:** Polygenic selection test results for the low $N_0$ model.

**Figure S4:** Permutation test results.

**Figure S5:** *Ne* estimates from simulations.

**Figure S6:** Histogram of mean *F*-values for each individual.

**Figure S7:** Schematic of data filtering.

**Figure S8:** Density of mean site depth.

**Figure S9:** Example of sliding–window analysis to detect over-assembled regions (OARs).

#### *Supplementary Tables*

**Table S1:** List of putative OARs.

**Table S2:** Cut-off values for filtering singletons.

### *Materials and Methods*

#### *Simulating Holstein tip ages*

Neutral genealogies were simulated using *msprime* (Kelleher *et al.*, 2016) to determine the mean tip length, and hence the background distribution of SDS in the absence of selection. We either simulated the Holstein population demography inferred by Boitard et al. (2016b), rounding estimated population sizes to the nearest integer, or with the present–day $N_e$ equal to 49 (Sørensen et al., 2005). We refer to these outputs as the 'High $N_0$' and 'Low $N_0$' models. 1,000 simulations were performed for each number of samples, ranging from 10 to 1,050. The mean tip length was calculated over all 1,000 simulations; 95% confidence intervals were calculated from 1,000 bootstraps. We fitted a linear model to the log of mean tip length against the log number of samples, and used it to predict the average tip age for 204 alleles, which is the number of diploid haplotypes used in the study. *B. taurus* are somewhat inbred (Sørensen *et al.*, 2005), which increases within–individual relatedness, and could reduce the number of unique alleles (see Nordborg and Donnelly (1997) for an example with self–fertilisation). Estimates of the inbreeding coefficient $F$ (Wright, 1951), which measure the reduction in heterozygosity, range from –0.15 to 0.35, with a mean of 0.059 (Figure S6; methods outlined below). Given this low mean value, we assumed two unique alleles per individual.

#### *Genome Data Extraction*

Whole genome sequencing for 102 Holstein bulls and cows were done by Illumina and BGI short-read paired-end sequencing in various laboratories with read lengths of 100 base pairs and up. No raw reads were shorter than 90 basepairs.

Bulls were selected for sequencing had high genetic representation in the present–day Holstein population. Sequencing of close relatives was avoided. Individuals were born between approximately 1980 and 2010. DNA was extracted from tissue, blood, or semen samples. Read data were processed according to the 1,000 Bull Genomes Project (Daetwyler *et al.*, 2014). Briefly, data were trimmed and quality filtered using Trimmomatic version 0.38 (Bolger *et al.*, 2014). Reads were aligned to the ARS-UCD-1.2 bovine genome assembly (Rosen *et al.*, 2018) (https://sites.ualberta.ca/~stothard/1000_bull_genomes/ARS-UCD1.2_Btau5.0.1Y.fa.gz) with the *B. taurus* Y chromosome assembly from BTau-5.0.1 added. Alignment was performed with bwa version 0.17 (Li & Durbin, 2009). Samtools (Li *et al.*, 2009) was used for sorting and marking of PCR duplicates. Base qualities were recalibrated using Genome Analysis Toolkit (GATK; McKenna et al. [2010]) version 3.8 using a set of known variable sites (Schnabel and Chamberlain, *unpubl*). GVCF files were formed using GATK's Haplotype Caller. Genotypes were called using GATK's GenotypeGVCFs.

##### *Initial filtering*

Figure S7 outlines a schematic of the data filtering. We first used *VCFtools* (Danecek *et al.*, 2011) to obtain a baseline list of biallelic SNPs. We removed indels and sites where the genotype was unknown in any individual. For each autosome, we obtained the mean depth for each remaining site using *VCFtools*' '--site-mean-depth' option. Figure S8 shows the depth distribution for these sites after initial filtering. We fitted a Poisson distribution to these data that had the same mean (9.76) as observed in the dataset. We determined the expected coverage range based on the 99.5% quantile range of the fitted distributions, which equalled 2 to 20. We subsequently removed sites that had mean coverage outside this range. We hence retained 6,873,371 of 20,010,175 initial variants (all entries in each autosome VCF file, including indels); we denote this dataset as the 'filtered' dataset.

##### *Finding putatively over–assembled regions*

Scaffolds of different genetic segments that each carry highly identical repeated regions can be 'over–assembled', where very similar chromosome regions were anchored to a single location (Chaisson *et al.*, 2015). These over–assembled regions (hereafter OARs) manifest themselves in aligned data as having high levels of heterozygosity, and elevated apparent coverage. If not corrected for, they can be misclassified as selected sites (e.g., subject to partial sweeps or balancing selection). We used a sliding window method to identify putative OARs in the dataset. For each chromosome, in each window we calculated (i) the number of sites where the reference allele had a frequency between 49% – 51%, (ii) the mean heterozygosity for each SNP (defined as the number of heterozygotes among the 102 individuals), and (iii) the mean summed allele depth. We used overlapping windows, each of size 500 SNPs with a step size of 10 SNPs. We first analysed all chromosomes to determine the distribution of values produced per window. We then re–ran the analyses, classifying windows as OARs if values for all three statistics belonged to the top 99.5% of their respective distributions, merging overlapping windows. We subtracted 1 from the start position of each region so that the leftmost boundary would also be excluded (if using '--bed-exclude' in *VCFtools*). Figure S9 shows an example where a region at the beginning of chromosome 1 was identified as an OAR. Overall, 5 OARs comprising 5,880 SNPs were identified (Table S1), which were subsequently masked from the rest of the pipeline.

##### *Calculating inbreeding coefficients*

Inbreeding coefficients were estimated using the '--het' option of *VCFtools*, which reports *F*-statistics for each chromosome per individual. Individual *F*-values (Figure S6) were calculated by taking the mean over all chromosomes, weighted by the chromosome size.

##### *Obtaining SDS analysis inputs*

***Test SNPs:*** Focal SNPs were those with an alternate allele frequency between 5% and 95%, and where each genotype was observed at least once amongst all samples. We then polarised values to determine the ancestral and derived states. Genome data, mostly from Wu *et al.* (2018), was downloaded from the NCBI SRA (https://www.ncbi.nlm.nih.gov/sra) for Bison (*Bison bison*, SRR6448737, SRR6448738, SRR6448739, SRR6448740), Wisent (*Bos bonasus*, SRR6448670, SRR6448682, SRR6448683, SRR6448684), Gaur (SRR6448732, SRR6448733, SRR6448734, SRR6448735), Gayal (*Bos frontalis*, SRR6448578, SRR6448580, SRR6448581), Banteng (*Bos javanicus*, SRR6448720 and SRR6448721), and as well as data for Yak (*Bos grunniens*, SRR5641601). Ancestral SNPs were determined by counting which allele was present in a majority of these related species; note that some SNPs for some species carried neither the reference nor the alternate alleles. We only retained SNPs where the called ancestral allele matched either the reference or alternate allele in our dataset. 3,405,023 SNPs were retained for testing.

***Singletons:*** Raw singleton data was extracted from the filtered Holstein dataset using *VCFtools*' '--singleton' option. This option identified both true singletons and private doubletons (i.e., where an allele is unique to an individual but present as a homozygote). Only true singletons were retained for analyses. To test whether a singleton had the same coverage as the non–singleton allele, we extracted the sequence depth for both alleles and retained sites satisfying the following criteria. First, the total allele depth was between 2 and 20 inclusive. Second, either (a) if the summed depth over both alleles exceeded 5, then the binomial probability of the observed allele depth exceeded 0.1; or (b) a stricter manual cut–off was applied if the total allele depth equalled 5 or less. Table S2 outlines the cut–off values used; 554,402 of 765,822 singletons were subsequently retained.

***Other parameters:*** The SDS method requires a 'singleton observability' probability, to predict how likely it is that a singleton will be detected by genome sequencing. Following Field et al. (2016) we used the mean depth per individual, obtained using the '--depth' option in *VCFtools.* It was also necessary to state the genetic boundaries between which analyses were carried out; we used a starting point of 1 and end points equal to the reported size of each autosome in *B. taurus*, as obtained from the ARS–UCD 1.2 genome assembly (https://www.ncbi.nlm.nih.gov/genome/?term=txid9913[orgn]).

Raw SDS values were calculated by fitting a gamma distribution to observed singleton distances, and comparing it to the expected distribution for the neutral case. We used the scripts provided by Field et al. (2016) (https://github.com/yairf/SDS) to generate the expected shape values for the gamma distribution for both the high and low $N_0$ models. Finally, we used a value of $10^{-7}$ to initiate the search for a maximum value in likelihood space.

##### *Calculating SDS scores and their significance for individual SNPs*

Out of 3,405,023 input SNPs, we retained and assigned scores to 1,877,770 of these. SDS scores were not assigned to a SNP if more than 5% of individuals did not harbour any singletons upstream or downstream of the SNP. This cut–off tended to exclude SNPs in telomeric regions. Furthermore, SDS scores were not calculated for chromosome 25 as an insufficient number of singletons were present across all individuals after data filtering. Raw SDS scores were standardized using 18 bins, based on alternate allele frequencies at the scored SNP (i.e., from 5% to 10%, from 10% to 15%, etc.). Standardised SDS scores (denoted sSDS) were obtained by subtracting the bin mean score from individual measures, and dividing by the bin standard deviation.

Statistical analyses were carried out in R (R Core Team, 2021). *P*–values for each SNP were determined by calculating the probability of the observed sSDS values from a standardised normal distribution. Significance was determined using a Bonferroni–corrected cut–off of 0.05/(1,877,770) $\approx 2.7 \times 10^{-8}$; note this cut-off is conservative as it does not account for potential linkage disequilibrium between SNPs (Moskvina & Schmidt, 2008). The false–discovery rate (FDR) of each SNP was calculated using the 'qvalue' package (Storey *et al.*, 2021); we highlighted SNPs with an FDR of less than 0.05.

*Data sources*

Version 104 of the GTF gene annotation file for the ARS–UCD 1.2 assembly was downloaded from Ensembl (available from https://www.ensembl.org/Bos_taurus/Info/Index). Bedtools v2.29.0 (Quinlan & Hall, 2010) was used to obtain genetic annotations 10kb up– and downstream of each Bonferroni–significant SNP; overlapping windows were merged.

Milk fat percentage and milk protein percentage QTLs were obtained from van den Berg et al. (2020). We retained only those QTLs for which effect sizes were reported in both Holstein breeds that were studied (French and German Holstein), and the effect was in the same direction in each. sSDS scores were then obtained for the nearest SNP to each QTL. Some QTLs lie close to the telomeres, where sSDS statistics were not available. These QTLs were not considered further; for each of the remaining QTLs, we identified the SNP nearest to it and assigned the sSDS value at that site to the QTL. Since QTL effect directions were given for the alternate allele (I. v. d. Berg, personal communication), we re-polarised sSDS scores for SNPs where the ancestral allele did not match the reference allele by switching its sign. We additionally re-polarised scores so that a positive value indicates selection acting on the trait-increasing allele (Field *et al.*, 2016). 30 of 138 milk fat QTLs had scores assigned to them, while 78 of 176 milk protein QTLs had scores assigned to them.

Stature QTLs were obtained from Bouwman et al. (2018), which identified 164 QTLs in several *B. taurus* breeds, including 7 Holstein populations from different countries. We initially extracted 114 QTLs for which an effect was inferred from at least 5 of 7 Holstein populations. Positions were given relative to the UMD 3.1

assembly; we subsequently extracted sequence 100bp up– and downstream of the position and remapped to ARS–UCD 1.0.25. 106 QTLs were re–aligned without gaps; of the remaining 8, 4 were located close to rearrangements and discarded, while the remaining 4 were kept. After removing those on chromosome 25, 107 QTLs were retained for downstream analysis. We analysed this full QTL set and a subset where effect sizes were reported in 6 of 7 Holstein breeds (containing 78 QTLs). We obtained the sSDS value of SNPs near QTLs and re-polarised them so that positive scores indicated selection on the trait-increasing allele. Overall, sSDS values were assigned to 58 QTLs with effect sizes in at least 5 of 7 Holstein populations, and 42 QTLs with effect sizes in at least 6 of 7 Holstein populations.

*Statistical analyses of polygenic selection*

To determine whether evidence exists for polygenic selection acting on QTLs underpinning these traits, we applied the 'cor.test' function in R to calculate the Spearman's rank correlation coefficient between sSDS values of the nearest SNP to the QTL, and the absolute $\log_{10}$ *p*-value of it as proxy for effect size. We also used the 'lm' function to determine and plot a linear regression between these values (see Figure 3 in the main text, S2, S3). To implement the permutation test, we subsampled SNPs from the genome that were from the same chromosome and derived-allele frequency bin as the SNPs used in the polygenic selection tests analyses; we also changed the sign of the sSDS value of the sampled SNP if this change was also made to the score associated with the corresponding true SNP. We then calculated and obtained the Spearman's rank coefficient between these permuted sSDS values and the QTL *p*-values. 1,000 subsampled datasets were used to generate the neutral distribution; significance was determined by normalising the true rho value (subtracting the mean and dividing by the standard deviation of permuted values), then calculating the two-tailed *P*–value from the standard normal distribution.

##### *Power simulations given Holstein demography*

To estimate the power of the SDS methods when applied to the Holstein breed, we implemented a simulation pipeline as used in previous studies of polygenic adaptation (Field *et al.*, 2016; Speidel *et al.*, 2019). We used the *simuPOP* program (Peng & Kimmel, 2005) to simulate the trajectory of either a neutral variant, or a partial selective sweep with homozygote selection coefficient of 0.2 (and a heterozygote value of half that value). The target mutation appeared *de novo* and we sampled individuals when it reached a frequency of 70%. We assumed the Holstein population demography as inferred by Boitard et al. (2016). Simulated trajectories were then piped into the coalescent simulation program *mbs* (Teshima & Innan, 2009) to generate polymorphism data for 204 haplotypes. We assumed a per-site recombination rate of $3.66 \times 10^{-9}$ (Boitard *et al.*, 2016), and a mutation rate of either $1.21 \times 10^{-8}$ as inferred from parent-offspring trios of dairy cattle (Harland *et al.*, 2018), or one ten-fold higher. A region of 10Mb long was simulated, with the target mutation appearing in the centre of it. Diploid genotypes were formed by randomly pairing haplotypes. SDS scores were calculated using the same pipeline and parameter values as for the data analyses. We also recorded the number of singletons per individual, and their distribution per simulation. 100 replicate simulations were run for each case. The confidence intervals for singleton distributions were calculated using

1,000 bootstraps. When plotting the distribution of SDS scores for the high-mutation case (Figure 5 in the main text), they were first normalised by subtracting the mean of the neutral case from each value, then dividing by the standard deviation of neutral scores.

To estimate the effective population size $N_e$ that would yield the simulated diversity, we use the statistic outlined by Watterson (1975) that relates an estimator of the population-level mutation rate to the number of segregating sites:

$$\theta = 4\ N_e\ \mu = \frac{S}{\sum_{i=i}^{n-i} 1/i} \quad (1)$$

where $\mu$ is the mutation-rate in the region; $S$ the number of segregating sites; and $n$ the number of samples (204 in our simulations). We hence estimated $N_e$ by obtaining the number of segregating sites, calculated the right-hand side of Equation 1, then divided by $4\mu^{*}L$ for $\mu^{*}$ the nucleotide mutation rate (1.21 x $10^{-8}$) and $L$ the length of the simulated genome (10Mb).

#### *References for Methods*


Berg, I. van den, Xiang, R., Jenko, J., Pausch, H., Boussaha, M., Schrooten, C., *et al.* (2020) Meta-analysis for milk fat and protein percentage using imputed sequence variant genotypes in 94,321 cattle from eight cattle breeds. *Genet. Sel. Evol.*, **52**, 37.

Boitard, S., Rodríguez, W., Jay, F., Mona, S. & Austerlitz, F. (2016) Inferring Population Size History from Large Samples of Genome-Wide Molecular Data - An Approximate Bayesian Computation Approach. *PLoS Genet.*, **12**, e1005877.

Bolger, A.M., Lohse, M. & Usadel, B. (2014) Trimmomatic: a flexible trimmer for Illumina sequence data. *Bioinformatics*, **30**, 2114–2120.

Bouwman, A.C., Daetwyler, H.D., Chamberlain, A.J., Ponce, C.H., Sargolzaei, M., Schenkel, F.S., *et al.* (2018) Meta-analysis of genome-wide association studies for cattle stature identifies common genes that regulate body size in mammals. *Nat. Genet.*, **50**, 362–367.

Chaisson, M.J.P., Wilson, R.K. & Eichler, E.E. (2015) Genetic variation and the *de novo* assembly of human genomes. *Nat. Rev. Genet.*, **16**, 627–640.

Daetwyler, H.D., Capitan, A., Pausch, H., Stothard, P., Binsbergen, R. van, Brøndum, R.F., *et al.* (2014) Whole-genome sequencing of 234 bulls facilitates mapping of monogenic and complex traits in cattle. *Nat. Genet.*, **46**, 858–865.

Danecek, P., Auton, A., Abecasis, G., Albers, C.A., Banks, E., DePristo, M.A., *et al.* (2011) The variant call format and VCFtools. *Bioinformatics*, **27**, 2156–2158.

Field, Y., Boyle, E.A., Telis, N., Gao, Z., Gaulton, K.J., Golan, D., *et al.* (2016) Detection of human adaptation during the past 2000 years. *Science*, **354**, 760–764.

Harland, C., Coppieters, W., Mullaart, E., Charlier, C. & Georges, M. (2018) *Rate of de novo mutation in dairy cattle and potential impact of reproductive technologies*. Proc. World Congr. Genet. Appl. to Livest. Prod.

Kelleher, J., Etheridge, A.M. & McVean, G. (2016) Efficient Coalescent Simulation and Genealogical Analysis for Large Sample Sizes. *PLoS Comput. Biol.*, **12**, e1004842.

Li, H. & Durbin, R. (2009) Fast and accurate short read alignment with Burrows–Wheeler transform. *Bioinformatics*, **25**, 1754–1760.

Li, H., Handsaker, B., Wysoker, A., Fennell, T., Ruan, J., Homer, N., *et al.* (2009) The Sequence Alignment/Map format and SAMtools. *Bioinformatics*, **25**, 2078–2079.

McKenna, A., Hanna, M., Banks, E., Sivachenko, A., Cibulskis, K., Kernytsky, A., *et al.* (2010) The Genome Analysis Toolkit: A MapReduce framework for analyzing next-generation DNA sequencing data. *Genome Res.*, **20**, 1297–1303.

Moskvina, V. & Schmidt, K.M. (2008) On multiple-testing correction in genome-wide association studies. *Genet. Epidemiol.*, **32**, 567–573.

Nordborg, M. & Donnelly, P. (1997) The Coalescent Process With Selfing. *Genetics*, **146**, 1185–1195.

Peng, B. & Kimmel, M. (2005) simuPOP: a forward-time population genetics simulation environment. *Bioinformatics*, **21**, 3686–3687.

Quinlan, A.R. & Hall, I.M. (2010) BEDTools: a flexible suite of utilities for comparing genomic features. *Bioinformatics*, **26**, 841–842.

R Core Team. (2021) *R: A Language and Environment for Statistical Computing*. R Foundation for Statistical Computing, Vienna, Austria.

Rosen, B., Bickhart, D.M., Schnabel, R.D., Koren, S., Elsik, C., Zimin, A.V., *et al.* (2018) Modernizing the Bovine Reference Genome Assembly. *Proc. World Congr. Genet. Appl. Livest. Prod.*, 11–16.
Sørensen, A.C., Sørensen, M.K. & Berg, P. (2005) Inbreeding in Danish Dairy Cattle Breeds. *J. Dairy Sci.*, **88**, 1865–1872.
Speidel, L., Forest, M., Shi, S. & Myers, S.R. (2019) A method for genome-wide genealogy estimation for thousands of samples. *Nat Genet.*, **51**, 1321–1329.
Storey, J.D., Bass, A.J., Dabney, A. & Robinson, D. (2021) qvalue: Q-value estimation for false discovery rate control [WWW Document]. URL http://github.com/StoreyLab/qvalue [accessed on 2021].
Teshima, K.M. & Innan, H. (2009) mbs: modifying Hudson's ms software to generate samples of DNA sequences with a biallelic site under selection. *BMC Bioinform.*, **10**, 166.
Watterson, G.A. (1975) On the number of segregating sites in genetical models without recombination. *Theor. Popul. Biol.*, **7**, 256–276.
Wright, S. (1951) The genetical structure of populations. *Ann. Eugen.*, **15**, 323–354.
Wu, D.-D., Ding, X.-D., Wang, S., Wójcik, J.M., Zhang, Y., Tokarska, M., *et al.* (2018) Pervasive introgression facilitated domestication and adaptation in the *Bos* species complex. *Nat. Ecol. Evol.*, **2**, 1139–1145.

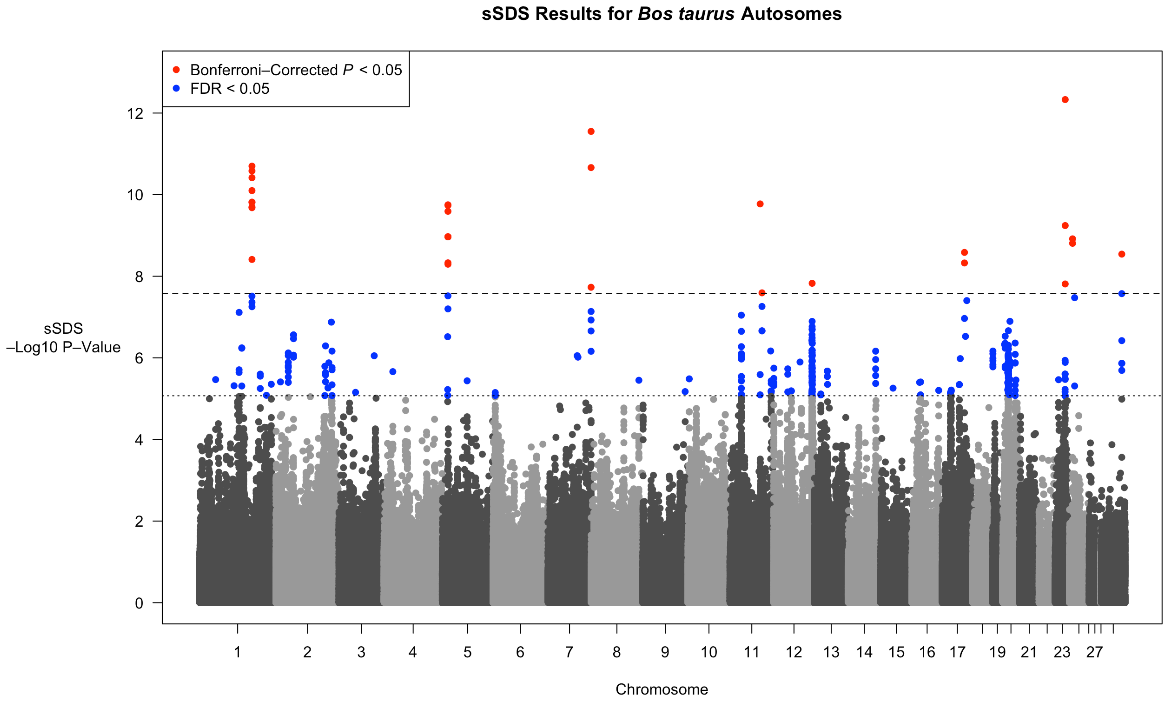


Figure S1: As Figure 2 in the main text, but assuming a low present–day $N_e$.

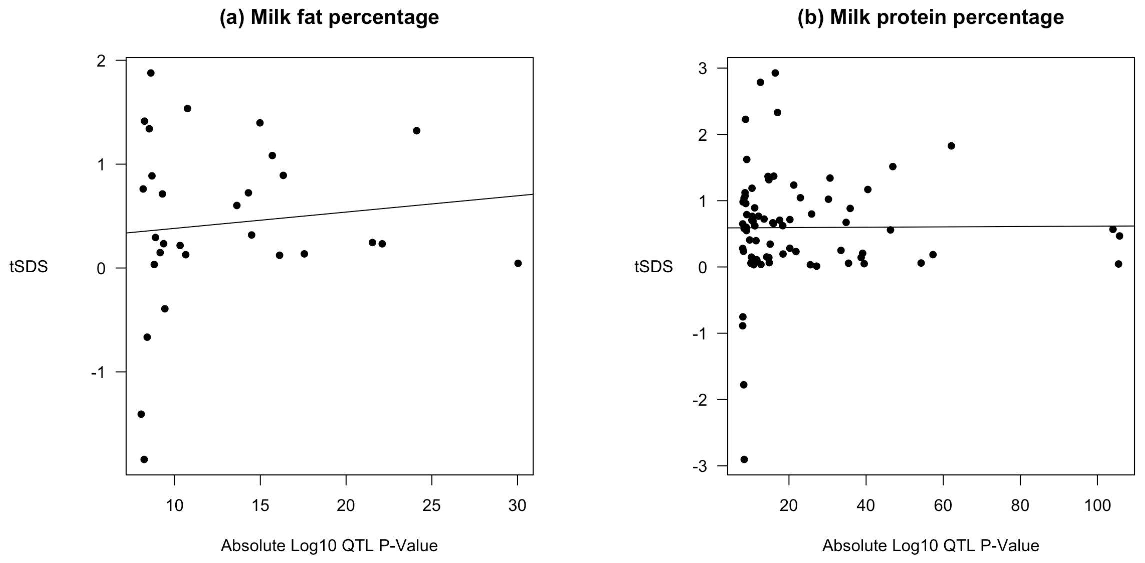


Figure S2: As Figure 3(a, b) in the main text but after removing an outlier point with a very low QTL *P*-value. *P*-values for correlations: milk fat percentage Spearman $\rho = 0.0818$, *P* = 0.673; milk protein percentage Spearman $\rho = 0.0255$, *P* = 0.826.

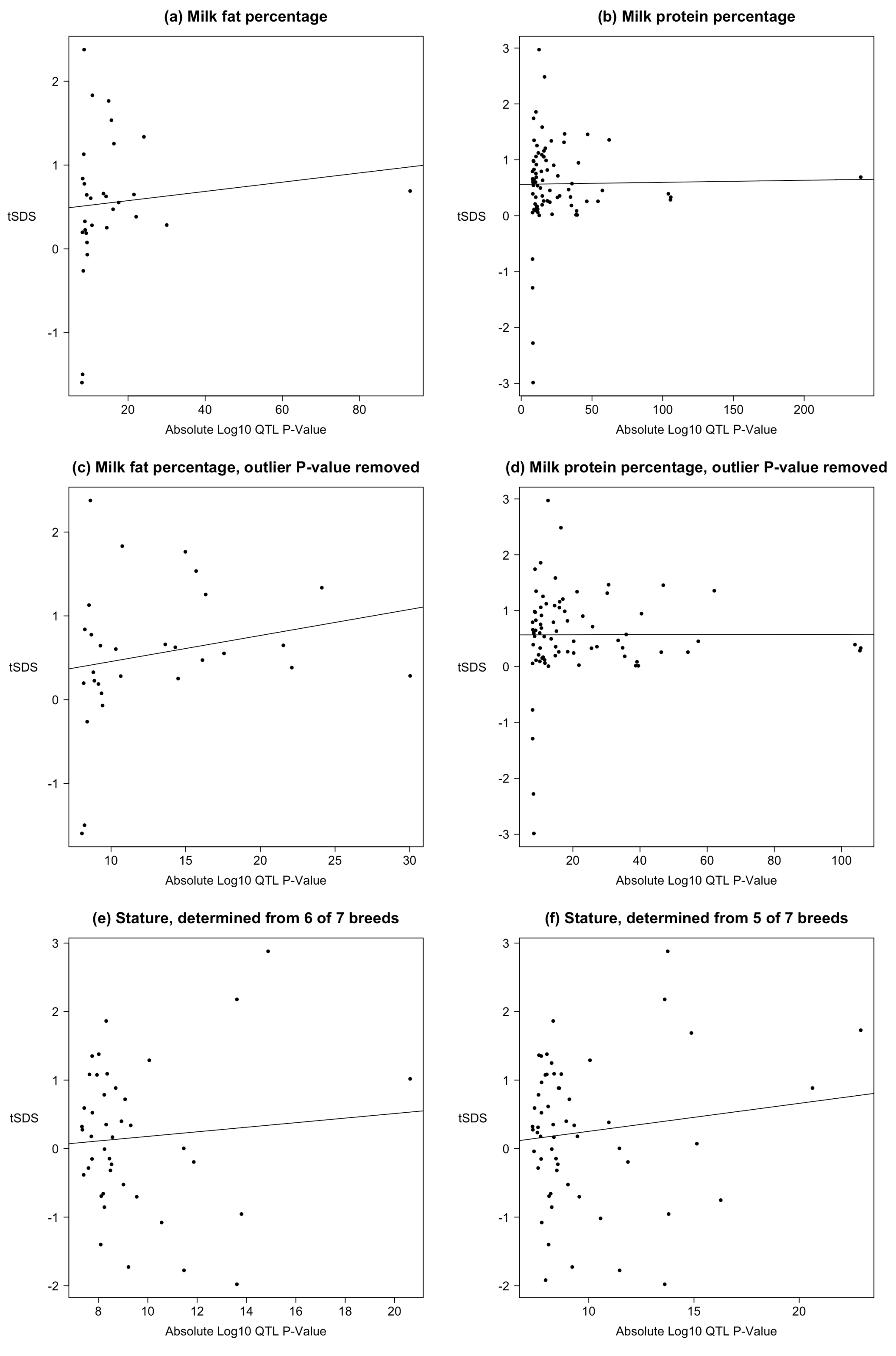


Figure S3: As Figure 3 in the main text and Figure S2, but assuming a low present–day $N_e$. $P$-values for correlations: (a) Spearman $\rho = 0.355$, $P = 0.0540$; (b) Spearman $\rho = 0.0515$, $P = 0.654$; (c) Spearman $\rho = 0.352$, $P = 0.0613$; (d) Spearman $\rho = 0.0446$, $P = 0.700$; (e) Spearman $\rho = -0.108$, $P = 0.498$; (f) Spearman $\rho = -0.0291$, $P = 0.828$.

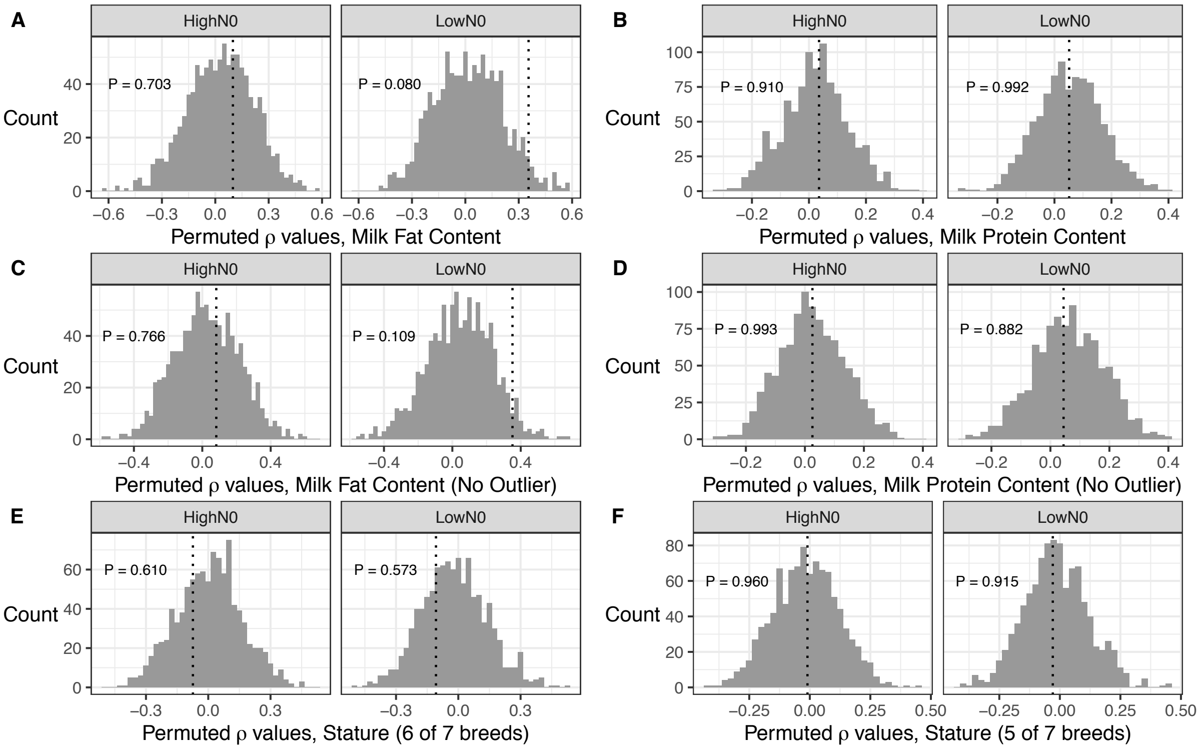


Figure S4: Histograms of Spearman's $\rho$ values for permuted datasets. Dashed line shows value from the true dataset; numbers in each subplot are *P*–values indicating to what extent the actual $\rho$ values lie outside those for the permuted datasets. Permutations are shown for (a) milk fat content, (b) milk protein content, (c) milk fat content with outlier point removed, (d) milk protein content with outlier point removed, (e) stature as inferred from 6 of 7 breeds, (f) stature inferred from 5 of 7 breeds.

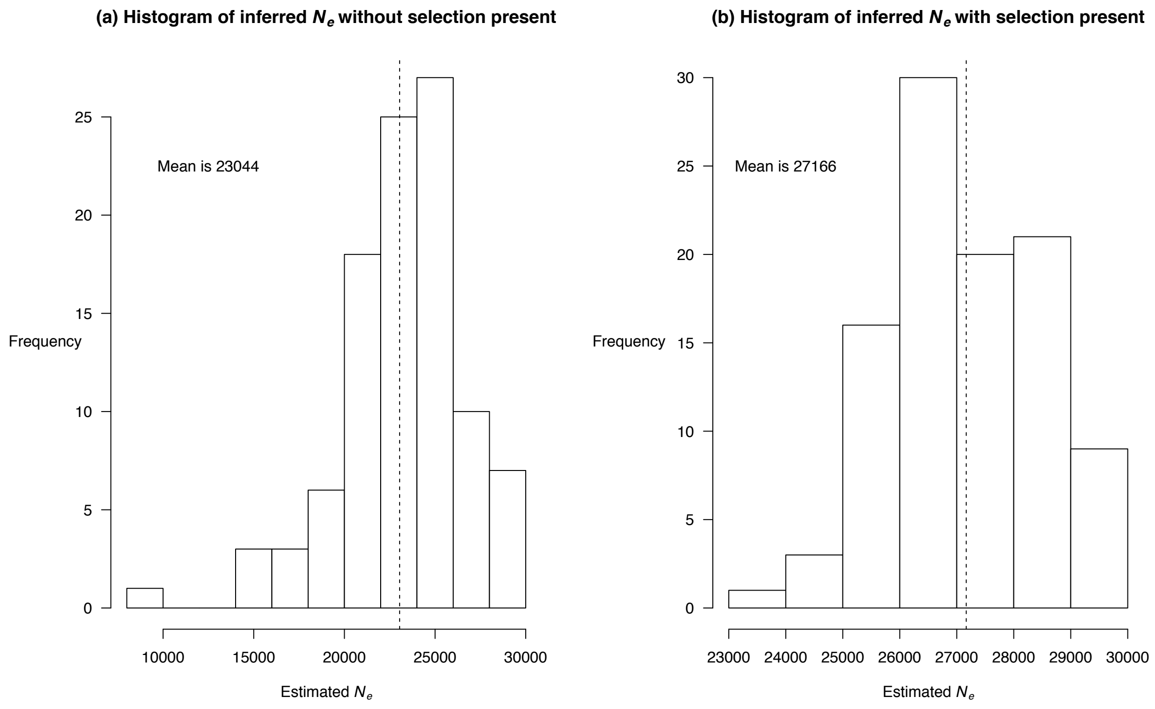


Figure S5: Estimates of a fixed *Ne* from simulations assuming Holstein demography, which would yield the same number of segregating sites. Vertical line shows the mean value, which is also listed in each subplot. Results are shown when the sweep was (a) absent or (b) present.

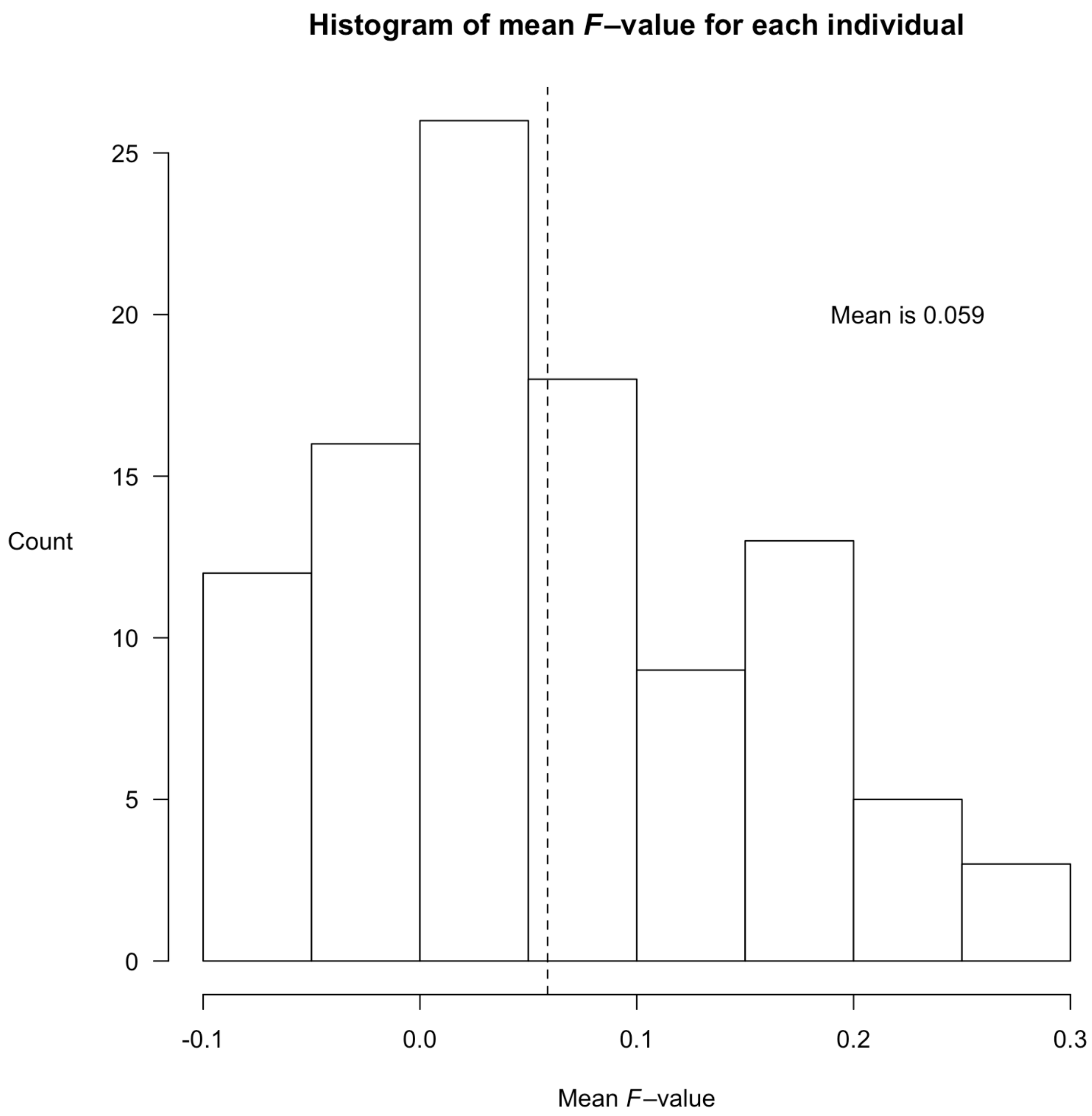


Figure S6: Histogram of mean inbreeding coefficient (*F*) for all individuals. Dashed line shows mean of mean values.

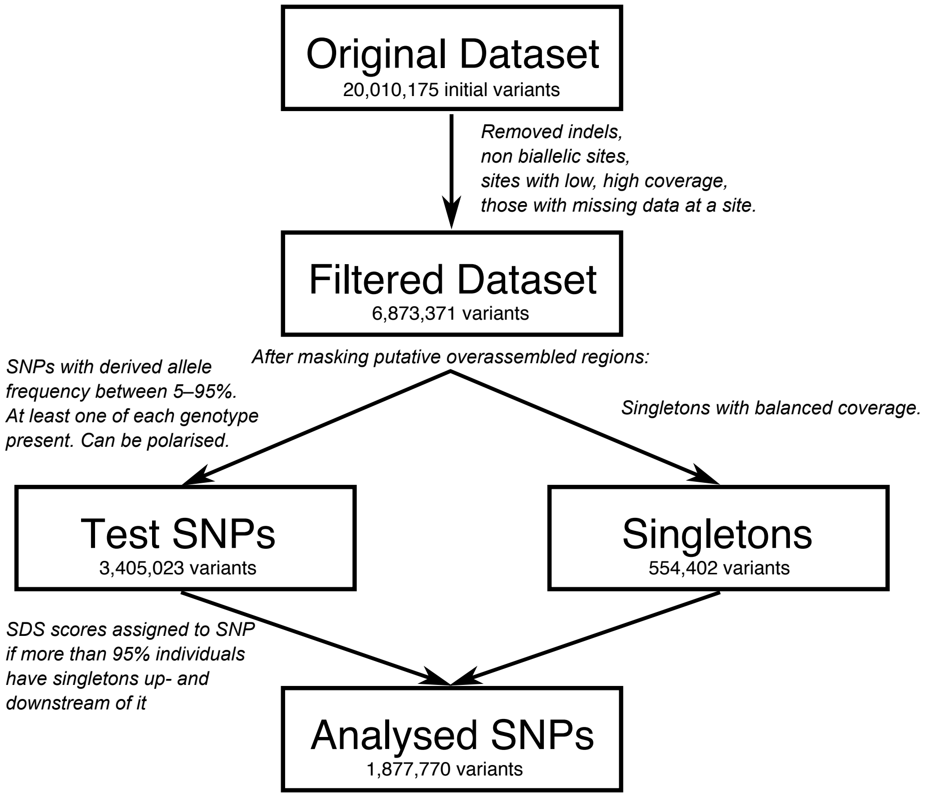


Figure S7: Schematic of data filtering.

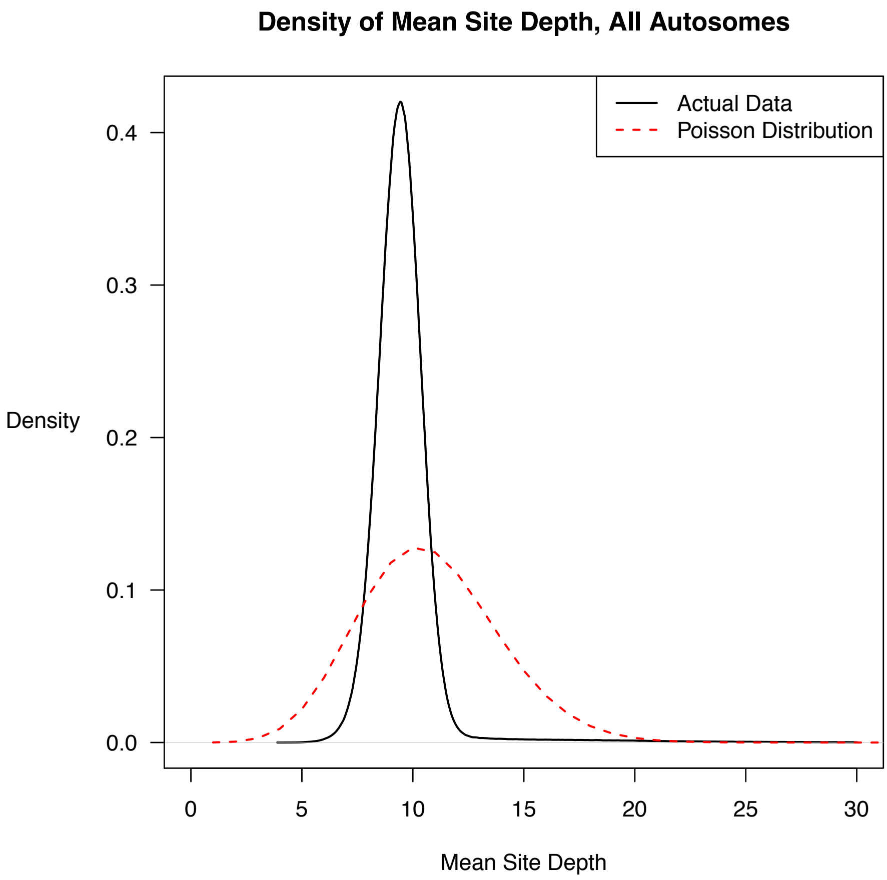


Figure S8: Density of mean site depth over all biallelic autosome SNPs before filtering. Note the plot only shows values between 0 to 30 for visual clarity; the maximum mean site depth is 1130.26. Red dashed line shows a Poisson distribution with the same mean (9.76) as the actual data.

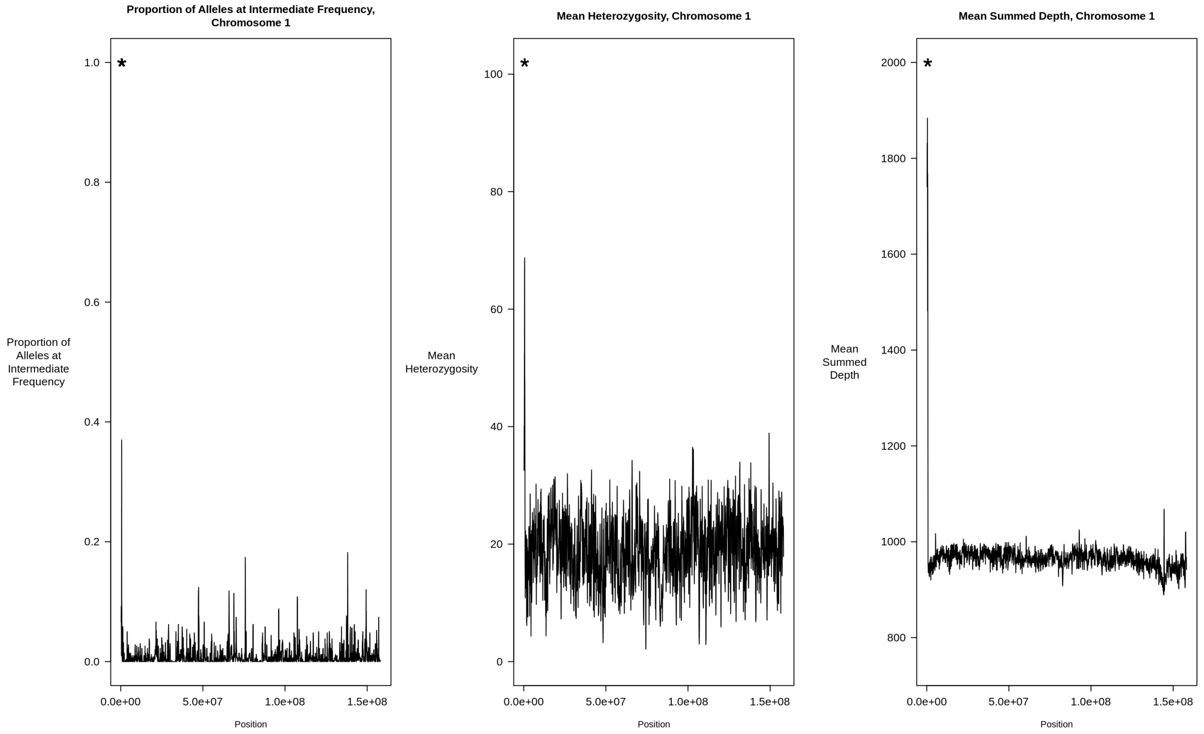


Figure S9: Example of the sliding window analyses to detect OARs for chromosome 1. The statistic being plotted is noted in the sub–window heading, as a function of the midpoint for each window. The asterisk denotes a detected OAR, between positions 481,018–637,494 (see Table S1 for a list of all putative OARs).

| Chromosome | Start Position | End Position | Number of SNPs in region |
|---|---|---|---|
| 1 | 481018 | 637494 | 1780 |
| 9 | 84666245 | 84704774 | 530 |
| 10 | 23439008 | 23622046 | 1840 |
| 13 | 10836694 | 10884908 | 720 |
| 15 | 49898125 | 50018198 | 1010 |

Table S1: List of putative OARs obtained for the *B. taurus* genomic dataset.

| If total allele depth equals: | Discard if one allele has less than this allele depth: |
|---|---|
| 2 | 1 |
| 3 | 1 |
| 4 | 2 |
| 5 | 2 |
| 6 | 2 |
| 7 | 2 |
| 8 | 2 |
| 9 | 3 |
| 10 | 3 |
| 11 | 4 |
| 12 | 4 |
| 13 | 5 |
| 14 | 5 |
| 15 | 6 |
| 16 | 6 |
| 17 | 7 |
| 18 | 7 |
| 19 | 8 |
| 20 | 8 |

Table S2: Table of cutoffs used to filter out singletons with unbalanced allele depths. Cutoffs for total allele depths 6–20 were determined by the minimum number of reads needed for a *P*–value of a binomial test (assuming equal allele coverage) to exceed 0.1. Cutoff for total allele depth 5 or less was set manually.
